## Supplemental information for "Molecular characterization reveals subclasses of 1q gain in intermediate-risk Wilms tumors"

Contents:

- Supplementary tables
- Supplementary figures

### Supplementary tables

**Table S1: Patient and sample characteristics.**

Labels and biomaterial identifiers of 30 patients with primary Wilms tumors, including clinical metadata, histopathological characterisation and quality control metrics.

IR: intermediate risk, HR: high-risk, c1a_pattern: chromosome 1 alteration pattern, qc: quality control, cov_med: median coverage, uq_reads: unique reads, qc_dup: percentage of duplicates.

**Table S2: Statistical tests to assess group differences in mutation burden**

**Table S3: Significantly differentially expressed genes associated with copy number gain and loss (Wilcoxon test, l2fc > +/- 0.138, FDR<0.2)**
l2fc: log2 fold change, FDR: false discovery rate.

**Table S4: Biological processes enriched in expression profiles (qvalue<0.05)**

**Table S5: Gene-level alteration overview, cancer genes only**
taf: tumor allele fraction, snv: single nucleotide variant, sv: structural variant, ctx: interchromosomal translocation, del: deletion, inv: inversion, dup: duplication, cna: copy number alteration, loh: cn neutral loss of heterozygosity, alteration: summary of the identified alterations (see methods for details).

Notation for SNVs: taf, protein_consequence, polyphen_label, sift_label. For SVs with breakpoints inside the gene: sv type, taf, position relative to the gene and sv size. For SVs that span the gene: sv type, taf and size. For SVs with breakpoints within 1 Mbp of the gene: sv type, taf, distance to the gene.

**Table S6: Details on somatic alterations affecting *CTNNB1, WT1* and *AMER1***
Summary of Table S5 with the same information and notation. c1a_pattern: chromosome 1 alteration pattern.

**Table S7: Expression profile genes that are both recurrently altered by CNs/SVs and recurrently overexpressed**
cna: copy number alteration, flag_sv_bp: has a structural variant breakpoint within 1 Mbp.

**Table S8: Chromosomal alterations per tumor**
cr l2fc: copy ratio log2 fold change, maf: minor allele fraction, estimated af: estimated tumor allele fraction, alteration: see methods for how chromosome arm level alterations are determined.

**Table S9: Somatic SVs**
sv: structural variant, ctx: interchromosomal translocation, del: deletion, inv: inversion, dup: duplication, tools: detection tools supporting the variant, sv_merged_coordinate: coordinate consensus by the tools, tumor_af_mean: mean tumor allele fraction of the tools, svlen: sv length, c1a_pattern: chromosome 1 alteration pattern.

**Table S10: Cancer genes on 1q overexpressed in 1q+ tumors (nzscore>1.98)**

### Figure S1. Mutation burden and correlations

**A)** Overview of alteration counts across tumors (rows) of various types (columns) with red lines depicting medians. Aneuploidy score: number of gain, loss or loss of heterozygosity (LOH) of full chromosomes and chromosome arms. Fraction of genome altered by copy number alterations (FGA). Non-synonymous single nucleotide variants (SNVs) and indels in coding genes. Number of somatic structural variants (SVs). **B)** SNV/indel burden correlates with age (*r=0.55* Pearson correlation) **C)** SNV/indel burden correlates with relative contribution of clock-like signature (SBS40, *r=0.66* Pearson correlation) **D)** Relative contribution of the mutational signatures we identified in this cohort: clock-like signature (SBS40), platinum-associated signature (SBS31), unknown signature (SBSA).

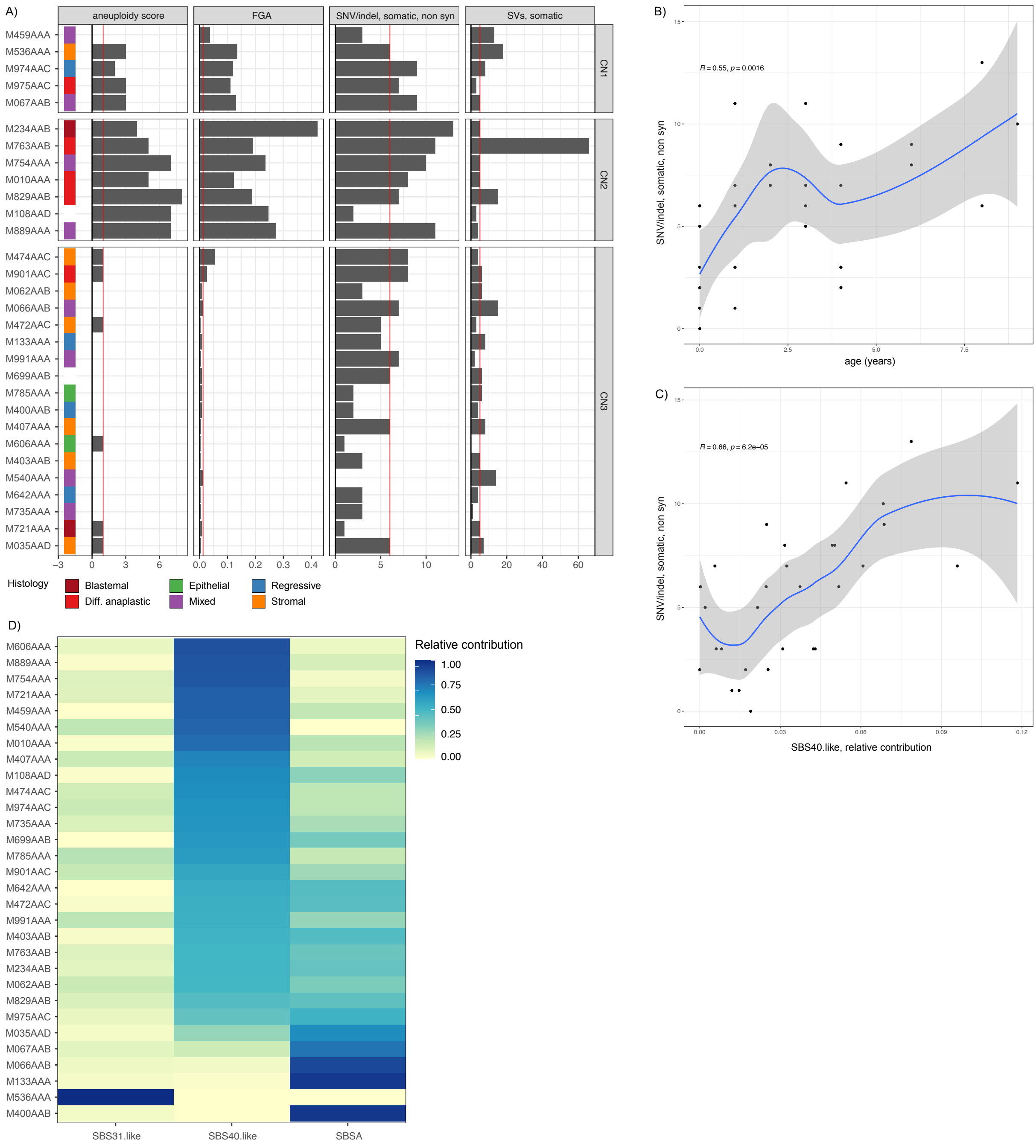

**Figure S2. Differentially expressed cancer genes associated with CN gain or loss**

Vulcano plot of differential gene expression results, comparing tumors with CN gain or loss with CN neutral tumors (l2fc > +/- 0.138, FDR<0.2). All genes are tested that are gained or lost in three or more tumors (recurrently altered). Significant cancer genes are labeled and coloured by location.

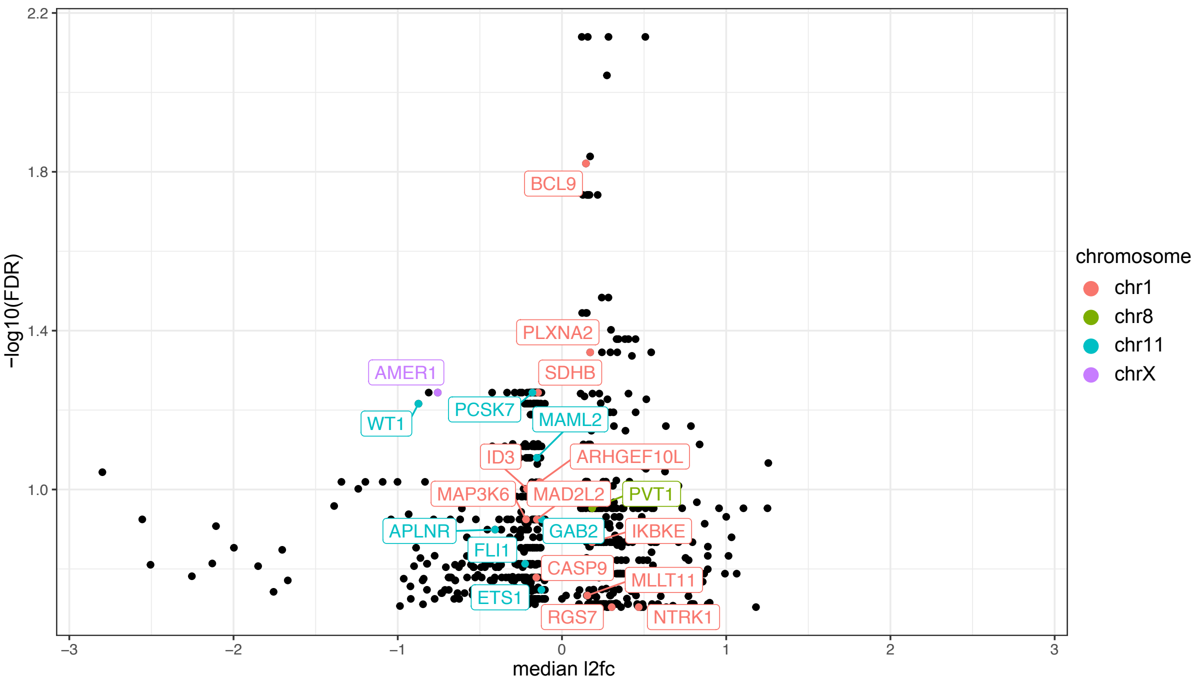

**Figure S3. Expression clusters**

UMAP of PCA 1-5 verifies the assignment of tumors to expression clusters based on their relative contribution of expression profiles. Tumors are annotated by their histological subtypes.

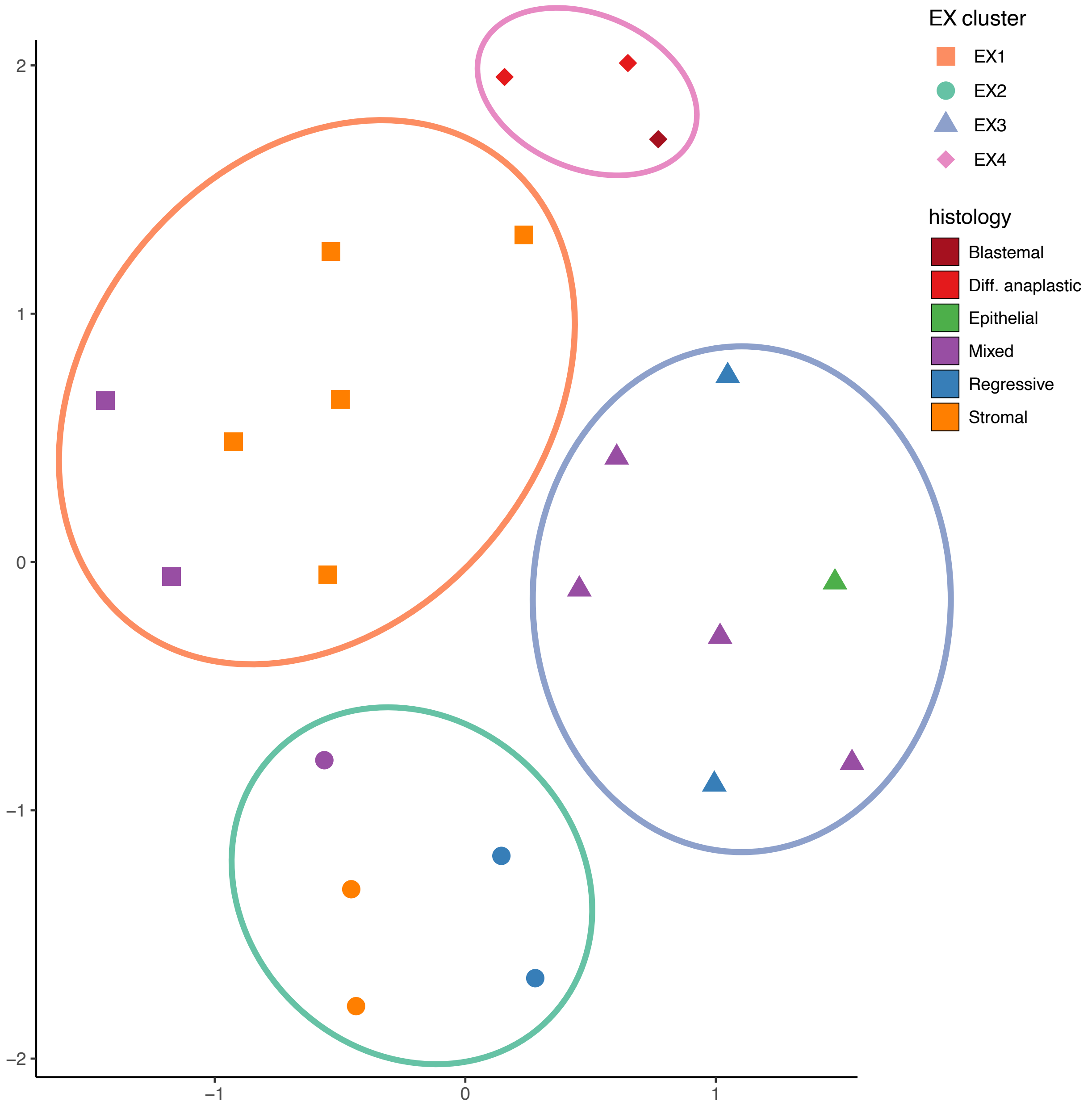

**Figure S4. Immune activating and suppressive genes**

To compare the immunological profile of tumors we selected cytokines and cell surface markers previously associated with activating or suppressive tumor microenvironment in Wilms tumor[1]. Although some tumors show high expression of activating *TNFA* or *IL6*, this is in conjunction with typical suppressive markers, e.g. *TGFB, IL10, IDO1/2* that are highly expressed in all EX2 tumors. Also the general markers of T-cell infiltration e.g. *CCL18* is upregulated in all EX2 tumors and among the top selected genes for the EX2 expression profile.

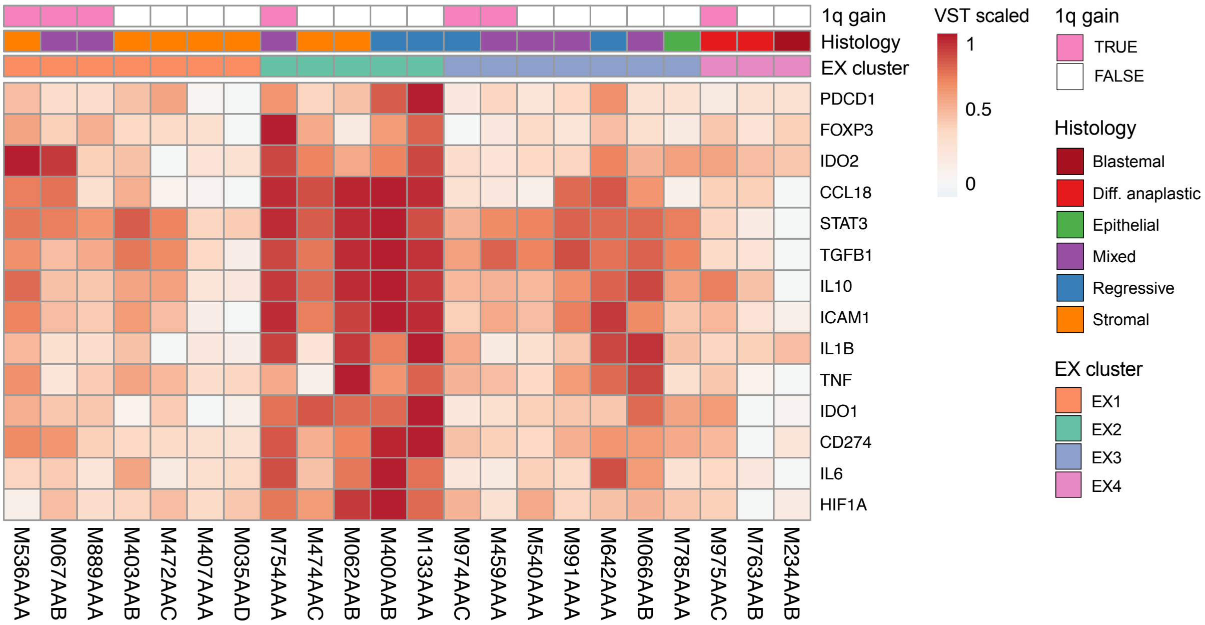


1. Systematic review of the immunological landscape of Wilms tumors (2021). Molecular Therapy - Oncolytics 22, 454–467.

### Figure S5. Somatic alterations in cancer genes of tumors

Full sized version of the Figure 2c oncoplot. Tumors are grouped by their expression cluster membership and annotated by their histological subtype. Genes included are all cancer genes (methods) and Wnt pathway genes. Alteration types included are: SNVs (purple) and SV breakpoints (green), as well as nearby or overlapping SVs (dark green) and CNs (orange) in case the tumor also has a significant change in gene expression (nzscore > +/-1.98). Genes are ordered by the number of tumors they are mutated in.

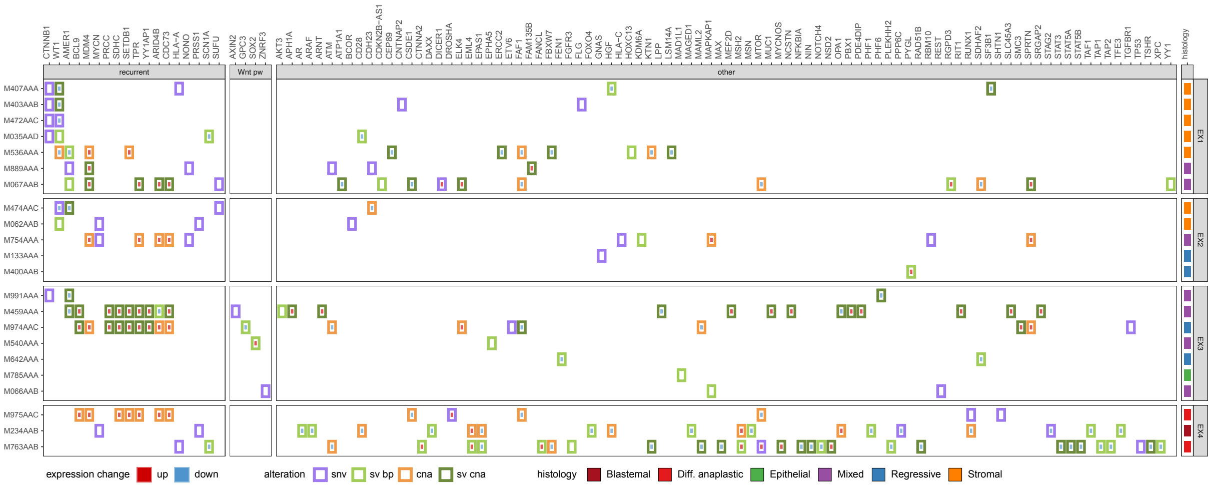

**Figure S6. Chromosomal alterations grouped by expression cluster**

Full sized version of Figure 2c. Genome-wide CN profiles of tumors grouped by their expression cluster membership with gains (red), losses (blue) and copy number neutral loss of heterozygosity (gold), overlain by SVs with translocations (CTX, black lines) and intrachromosomal variants (gray). Tumors are annotated by their histological subtype.

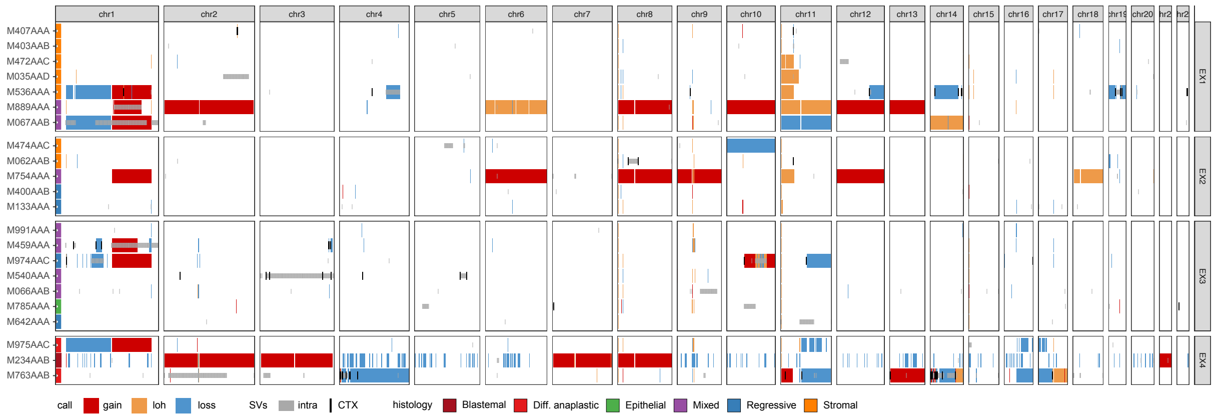

**Figure S7. Linking gene expression changes to recurrent CNs/SVs**

Expression clusters are characterized by increased expression of the expression profile gene sets in the tumors assigned to the cluster relative to the remainder of the cohort. To investigate the role of CNs/SVs in mediating these expression patterns, we assessed whether genes were recurrently upregulated in two or more tumors of the same expression cluster (>1.5 nzscore, 2498 genes) and/or have recurrent underlying CN gains or nearby SV breakpoints within 1 Mbp (272 genes). Comparison of these sets shows that 51 genes show recurrent upregulation and are also recurrently altered by CNs/SVs.

Overall, 2% of recurrently upregulated genes has a recurrent CN/SV (51/2498). 19 % of the recurrent CN/SV altered genes is upregulated in 2 or more tumors (51/272) And 23% of recurrently overexpressed genes has a CN/SV in at least one tumor (581/2498).

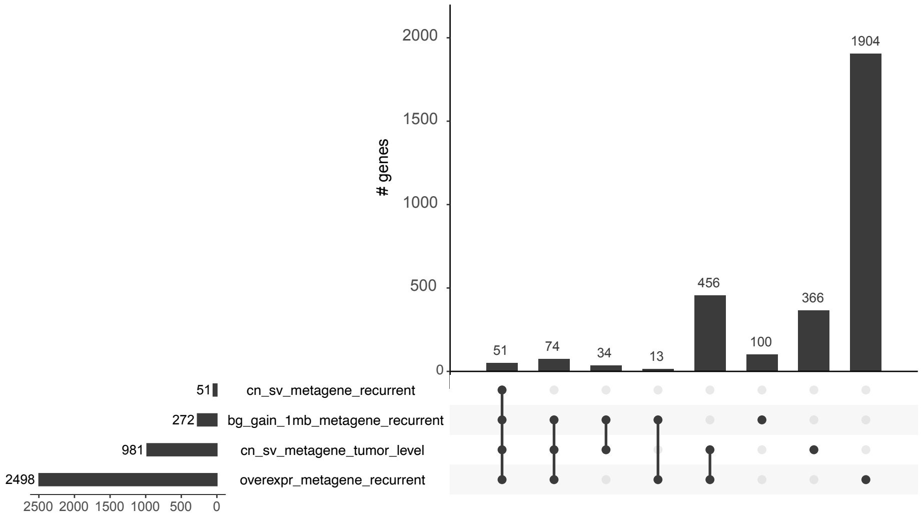

**Figure S8. Location of SVs and differentially expressed genes in 1q+ tumors**

To investigate the relationship between SVs and up- or downregulated genes on chromosome 1 (nzscore > +/- 1.98), we displayed them together in one figure. Genes are depicted as black vertical lines, cancer genes altered in two or more tumors are labeled. SVs are colored by type: duplications (DUP, red), deletions (DEL, blue), inversions (INV light purple), translocations, (CTX, dark purple lines). Distributions of over/under expressed genes are colored by the tumor's expression cluster (EX1, orange, EX2 green, EX3 blue, EX4 pink).

**A)** The EX2 tumors with focal 1p loss due to underlying SVs (M459AAA and M974AAC) downregulate genes nearby their breakpoints, whilst the tumors with full arm 1p loss show downregulation of genes across the chromosome arm. **B)** 1q+ tumors differ in the mutational mechanism of 1q gain (Figure 5a) and the genes that are overexpressed on 1q (Figure 5b), but we did not observe a relationship between the SVs and location of overexpressed genes. EX3 tumors overexpress most cancer genes (16 and 17 genes) and share seven genes (*BCL9*, *PRCC, SDHC, CDC73, SETDB1, TPR, YY1AP1*) whilst EX1 tumors share only *MDM4* overexpression.

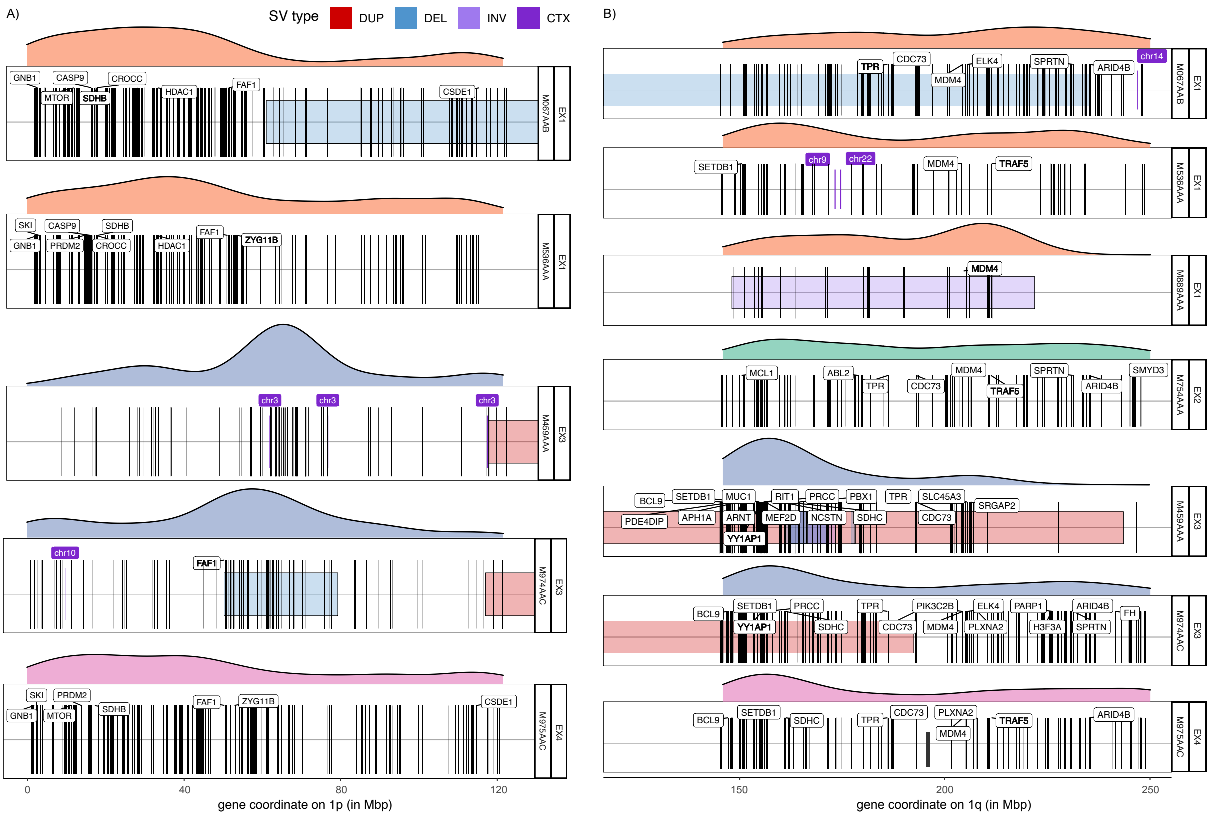
